# Highly expressed maize pollen genes display coordinated expression with neighboring transposable elements and contribute to pollen fitness

**DOI:** 10.1101/796060

**Authors:** Cedar Warman, Kaushik Panda, Zuzana Vejlupkova, Sam Hokin, Erica Unger-Wallace, Rex A Cole, Antony M Chettoor, Duo Jiang, Erik Vollbrecht, Matthew MS Evans, R Keith Slotkin, John E Fowler

## Abstract

In flowering plants, the haploid male gametophyte (pollen) is essential for sperm delivery, double fertilization, and subsequent initiation of seed development. Pollen also undergoes dynamic epigenetic regulation of expression from transposable elements (TEs), but how this process interacts with gene regulation and function is not clearly understood. To identify components of these processes, we quantified transcript levels in four male reproductive stages of maize (tassel primordia, microspores, mature pollen, and isolated sperm cells) via RNA-seq. We found that, in contrast to Arabidopsis TE expression in pollen, TE transcripts in maize accumulate as early as the microspore stage and are also present in sperm cells. Intriguingly, coordinated expression was observed between the most highly expressed protein-coding genes and neighboring TEs, specifically in both mature pollen and sperm cells. To test the hypothesis that such elevated expression correlates with functional relevance, we measured the fitness cost (male-specific transmission defect) of GFP-tagged exon insertion mutations in over 50 genes highly expressed in pollen vegetative cell, sperm cell, or seedling (as a sporophytic control). Insertions in genes highly expressed only in seedling or primarily in sperm cells (with one exception) exhibited no difference from the expected 1:1 transmission ratio. In contrast, insertions in over 20% of vegetative cell genes were associated with significant reductions in fitness, showing a positive correlation of transcript level with non-Mendelian segregation. The *gamete expressed2* (*gex2*) gene was the single sperm cell gene associated with reduced transmission when mutant (<35% for two independent insertions), and also triggered seed defects when crossed as a male, supporting a role for *gex2* in double fertilization. Overall, our study demonstrates a developmentally programmed and coordinated transcriptional activation of TEs and genes, and further identifies maize pollen as a model in which transcriptomic data have predictive value for quantitative phenotypes.

**Author Summary:** In flowering plants, pollen is essential for delivering sperm cells to the egg and central cell for double fertilization, initiating the process of seed development. In plants with abundant pollen like maize, this process can be highly competitive. In an added layer of complexity, growing evidence indicates expression of transposable elements (TEs) is more dynamic in pollen than in other plant tissues. How these elements impact pollen function and gene regulation is not well understood. We used transcriptional profiling to generate a framework for both detailed analysis of TE expression and quantitative assessment of gene function during maize pollen development. TEs are expressed early and persist, many showing coordinate activation with highly-expressed neighboring genes in the pollen vegetative cell and sperm cells. Measuring fitness costs for a set of over 50 mutations indicates a correlation between elevated transcript level and gene function in the vegetative cell. Finally, we establish a role in fertilization for the *gamete expressed2* (*gex2*) gene, identified based on its specific expression in sperm cells. These results highlight maize pollen as a powerful model for investigating the developmental interplay of TEs and genes, as well as for measuring fitness contributions of specific genes.

## Introduction

Sexual reproduction enables the segregation and recombination of genetic material, which increases genetic diversity in populations and contributes to the vast diversity of eukaryotes. In flowering plants, sexual reproduction requires the development of reduced, haploid gametophytes from sporophytic, diploid parents. The mature female gametophyte, the embryo sac, includes the binucleate central cell and the egg cell (reviewed in [1,2]), each of which is fertilized by a sperm cell to generate the triploid endosperm and diploid embryo, respectively. The mature male gametophyte, pollen, consists of a vegetative cell harboring two sperm cells (reviewed in [3,4]). In maize, male gametophytes arise from microspore mother cells in the tassel primordium. The transition from diploid sporophyte to haploid gametophyte occurs when these cells undergo meiosis, each resulting in four haploid microspores. Each microspore then undergoes two rounds of mitosis to produce the pollen grain, first generating the large vegetative cell and a smaller generative cell via asymmetric division, and then producing the two sperm from the generative cell. After the arrival of the pollen grain on the floral stigma, the vegetative cell transports the two sperm cells to the female gametophyte via pollen tube growth (reviewed in [5,6]). Accurate navigation of the pollen tube as it grows down the style is dependent on the architecture of the style’s transmitting tract [7] and possible additional signaling and recognition mechanisms that are poorly understood [8]. The final stages of pollen tube growth depend on a complex interplay of signals to guide the pollen tube to the ovule (reviewed in [9]).

In maize, a pollen tube must grow up to 30 cm through the silk to reach the female gametophyte, often competing with multiple pollen tubes to eventually enter the embryo sac and release its sperm cells for fertilization (reviewed in [2,5]). Across the plant kingdom, this competitive context for pollen tube development differs, depending on the pollen population as well as sporophytic characters (reviewed in [10]). In a highly competitive environment, successful fertilization is likely enhanced by pollen tubes functioning at full capacity [11–13], as generally only the first tube to reach the micropyle is permitted to enter the female gametophyte. The mechanisms preventing entry of multiple pollen tubes, known as the polytubey block, are not well-understood, but presumably act to reduce polyspermy, which typically leads to sterile offspring [6]. In Arabidopsis, mutant-associated fertilization of only the egg cell or the central cell does not fully activate the block, allowing for the attraction of an additional pollen tube and the completion of double fertilization with sperm cells from two different pollen tubes [14], a process known as heterofertilization. Heterofertilization occurs at low frequencies (0.5-5%, depending on the genetic background) in maize [15,16] and is thought to be associated with abnormal fertilization, but the details remain unclear. In maize, mutations in the genes *MATRILINEAL*/*NLD*/*ZmPLA1* and *ZmDMP* have been linked to pollen-induced production of haploid embryos and other seed defects, which are likely associated with aberrant events at fertilization [17–20] or soon after [21]. Thus, many mechanisms associated with both pollen tube growth and fertilization remain enigmatic.

Given their specialized biological functions and well-defined developmental stages, gametophytes are prime targets for transcriptome analysis. Initial studies of plant gametophytic transcriptomes in Arabidopsis pollen [22,23] and embryo sacs [24,25] described a limited and specialized set of transcripts and identified numerous candidate genes for gametophytic function. In maize, the first RNA-seq study of male and female gametophyte transcriptomes (mature pollen and embryo sacs) similarly identified subsets of developmentally specific genes, with pollen showing the most specialized transcriptome, relative to other tissues assessed [26]. More recently, RNA-seq has been carried out on additional stages of maize reproductive development, including pre-meiotic and meiotic anther cells [27–29], as well as sperm cells, egg cells, and early stages of zygotic development [30].

Gametophytic tissues are known to show dynamic expression of transposable elements (TEs). In Arabidopsis, global TE expression is derepressed at the late stages of pollen development, occurring in the pollen vegetative nucleus only after pollen mitosis II [31]. The pollen vegetative nucleus undergoes a programmed loss of heterochromatin, resulting in TE activation, TE transposition, TE small interfering RNA production, and subsequent increased RNA-directed DNA methylation [31–35]. A variety of functions have been ascribed to this male gametophytic “developmental relaxation of TE silencing” (DRTS) event [36], including the generation of TE small interfering RNAs that are mobilized to the sperm cells [37], and control of imprinted gene expression after fertilization [38]. However, the dynamics of TE expression during gametophytic development in a transposable element-rich species such as maize have not been investigated.

To provide a more full description of transcriptome dynamics across maize male reproductive development, including TE transcriptional activity, we generated RNA-seq datasets from tassel primordia, microspores, mature pollen, and isolated sperm cells. Using these data, we describe differential expression patterns of genes and TEs across these stages, uncovering a coordinated regulation of TEs and their neighboring genes in pollen grains. We then conducted a functional validation of such highly expressed genes by testing over fifty insertional mutations for male-specific fitness effects. Finally, these transcriptome data guided the discovery of mutant alleles in the sperm cell-enriched *gex2*, which induces seed development defects when present in the pollen parent, implying a role in fertilization.

## Results

### Experimental design and gene expression during maize male reproductive development

RNA-seq was performed on four tissues representing integral stages in maize male gametophyte development: immature tassel primordia (TP), isolated unicellular microspores (MS), mature pollen (MP), and isolated sperm cells (SC) (Fig 1A). Techniques were developed to efficiently isolate RNA from TP, MS, and SC (see Methods). RNA was extracted from the inbred maize line B73, with four biological replicates for each tissue. In addition, a single RNA replicate was isolated for the bicellular stage of pollen development (MS-B). Libraries were sequenced using Illumina sequencing (100 bp paired-end reads) and mapped to the B73 AGPv4 reference genome [39]. Principal Component Analysis (PCA) showed samples from each tissue clustering together along PC1 and PC2, which together explained 49.8% of the variance between samples (Fig 1B). One sample, SC1, had significant levels of ribosomal RNA (rRNA) contamination, as well as the fewest number of mapped reads (approximately 1 million). However, to maintain a balanced experimental design with a consistent false discovery rate (FDR), we chose to include SC1 in our analysis of gene expression patterns.

**Fig 1.**
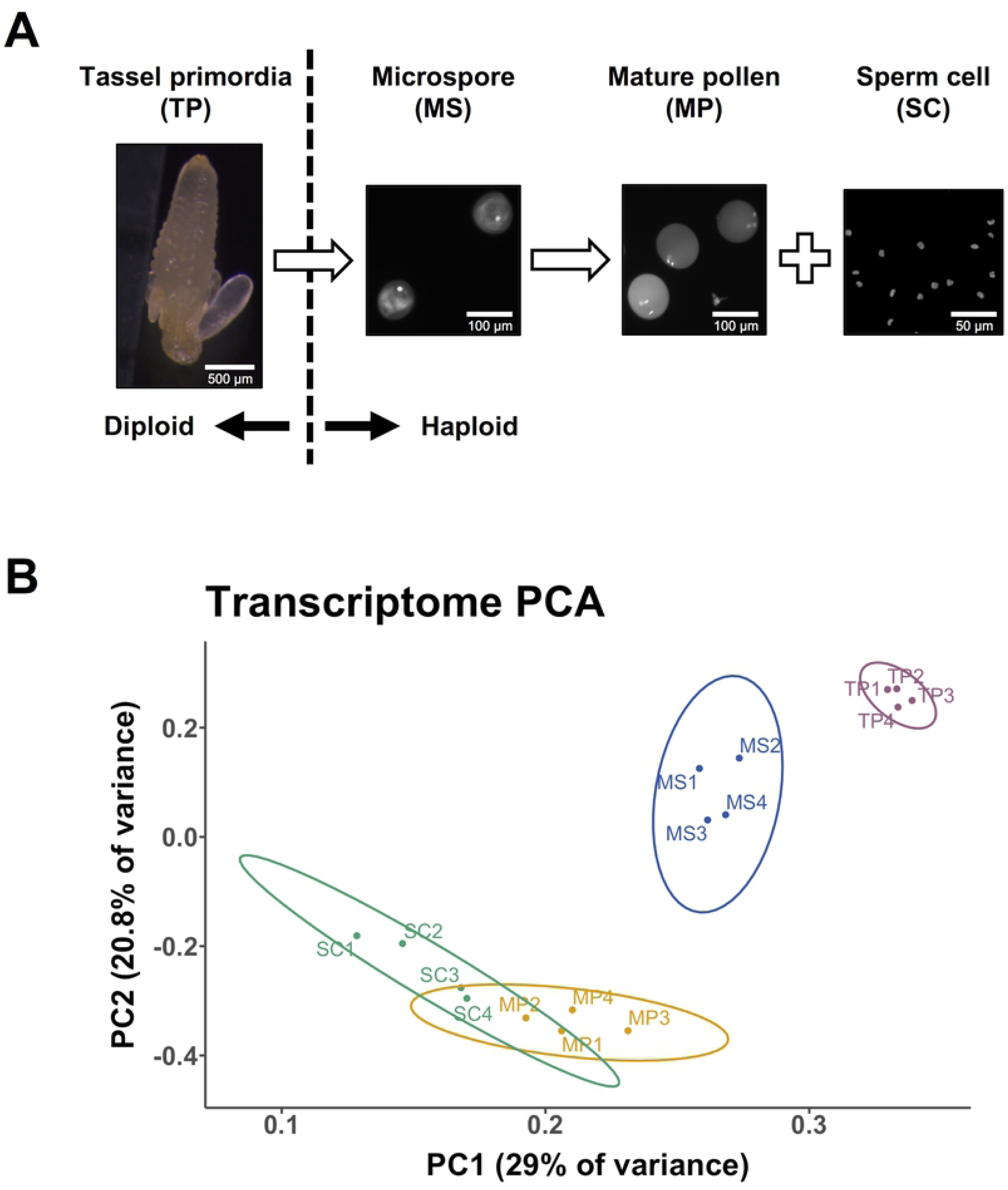
Experimental design and transcriptome replicate assessment. **(A)** mRNA was isolated from four developmental stages of maize male reproductive development, with four biological replicates for each: pre-meiotic tassel primordia (TP), post-meiotic, unicellular microspores (MS), mature pollen (MP), and sperm cells (SC). A single biological replicate of mRNA from the bicellular stage of pollen development was also isolated and sequenced (MS-B, not shown). Nuclei were stained with either DAPI or Dyecycle green. **(B)** Principal component analysis of genic transcriptomic data generated by this study, showing the 2 major components (explaining 49.8% of the variance) on x- and y-axis. The four biological replicates of each of the four sequenced tissues clustered with other replicates from the same tissue. TP and MS were clearly separated in principal component space, whereas SC and MP samples displayed less separation from each other.

Differential gene expression was defined in two ways: in the first, gene expression in later developmental stages was compared to the premeiotic, diploid tassel primordia (TP vs MS, TP vs MP, and TP vs SC); in the second, gene expression was compared between all adjacent developmental stages (TP vs MS, MS vs MP, MS vs SC, MP vs SC) (S2 and S11 Tables). Enriched GO terms highlighted the differences in gene expression among developmental stages and suggested consistency with the established functions of each tissue [22,26,30]. GO terms in MS were consistent with a post-meiotic tissue still at an early stage of development, with terms related to protein synthesis and transport, morphogenesis, and reproduction showing enrichment. MP showed more specific enriched GO terms, including those related to pollen tube growth, signaling, and actin filament-based movement. SC shared many GO terms with MP when compared to MS, but was uniquely enriched for GO terms related to epigenetic regulation of gene expression, such as gene silencing by RNA and histone H3-K9 demethylation.

### A subset of transposable elements in the maize genome show developmentally dynamic expression

To obtain a broad view of TE expression throughout maize development, the RNA-seq data for maize male reproductive development generated by our sequencing (samples with asterisks, S1 Fig) was combined with publicly available RNA-seq of nine-day old above-ground seedlings, juvenile leaves, ovules, another set of independently isolated sperm cells, and three independent studies of pollen RNA-seq [26,30,40,41] & SRP067853. The complete list of samples, their sequencing statistics, references and data availability can be found in S3 Table. All of the raw data were remapped using the same parameters (see Methods). Principal component analysis demonstrates that replicates of the same tissue and growth state typically group together (S1 Fig).

We aimed to identify the set of dynamically expressed TEs within the tissues sampled. Thus, the RNA-seq samples were used to calculate expression levels for each individual TE in the genome located more than 2 kb away from annotated non-TE genes. Our rationale was to avoid false positive signals of TE expression due to a TE residing within a gene, and to minimize the influence of read-through transcription from a nearby gene, which could not be distinguished from TE-initiated transcription. To relate TE expression comparatively across development, we used seedling tissue as a baseline against which other tissues were measured. Seedling was chosen for several reasons: it is not a reproductive tissue, it has low to average levels of TE expression, and a large number of TEs show no evidence of expression in this tissue (S2 Fig).

Apart from 18.3% of the annotated TEs that are near genes and analyzed separately (see below), we calculated the number of TEs with statistically significant expression differences in each tissue compared to the seedling reference. This identified the subset of TEs that are developmentally dynamic, meaning that they show differential expression in at least one tissue in our dataset compared to the seedling reference. Only 4.4% of all maize annotated TEs are developmentally dynamic, whereas 22.2% of TEs have detectable expression, but do not change in our dataset and therefore are developmentally static (Fig 2A). Finally, the majority of annotated TEs (55%) were not assessed in this analysis, either because no expression was detected in any dataset, or because their sequence lacks the polymorphisms necessary for mapping to a specific TE.

**Fig 2.**
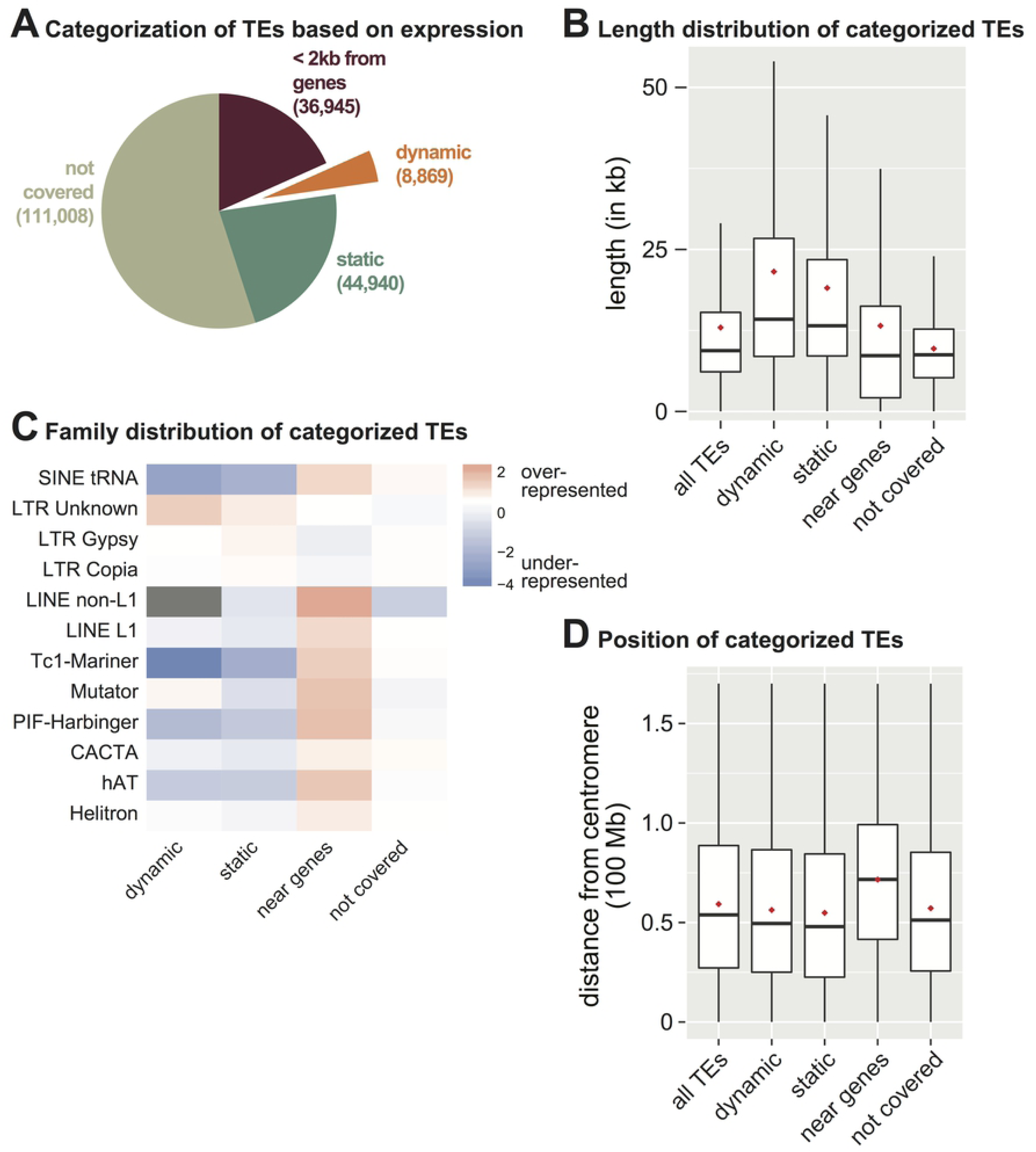
Characterization of developmentally dynamic transcription from transposable elements (TEs). **(A)** Distribution of different categories of TEs based on their expression. Number of TEs are in parentheses. **(B)** Length of TEs in the different TE categories from part A. The box represents lower and upper quartile, the line is the median, and the whiskers represent 10-90% range. Red asterisk denotes the mean. **(C)** Observed / expected Log2 ratios of TE family proportions in the different TE categories from part A. Grey indicates no data. **(D)** Distance from the annotated centromere for different TE categories from panel A.

Each TE category (Fig 2A) was interrogated for feature overrepresentation. Both dynamic and static TEs are longer than the genome average, and longer than the sets of TEs ‘not covered’ or ‘near genes’ (Fig 2B). The finding that expressed TEs as a group (dynamic + static) are longer correlates with Arabidopsis data where longer TE transcripts are overrepresented and differentially regulated when epigenetic repression is lost [42]. Expressed TEs show an under-representation for DNA transposon and SINE families, which are mainly within the ‘near genes’ set (Fig 2C). In contrast, the ‘LTR unknown’ TE annotation is over-represented in the dynamic TE set (Fig 2C). Since some LTR retrotransposons are enriched in the pericentromere [43], we tested if the dynamic TE set is enriched in the pericentromere compared to the genome average, but did not detect any correlation (Fig 2D). Therefore, we conclude that expressed TEs are generally longer elements, and the subset of developmentally dynamic TEs are enriched for uncharacterized LTR retrotransposons located throughout the genome.

### Transposable element transcript levels are up-regulated in the post-meiotic male reproductive lineage

From the developmentally dynamic TE set, we calculated the number of differentially expressed TEs in each tissue/stage compared to the seedling reference. In some tissues, such as tassel primordia and ovules, we observed a similar number of TEs up-regulated and down-regulated (Fig 3A), demonstrating that while there are shifts in which TEs are expressed, a genome-scale change in TE expression does not occur. In other tissues, such as juvenile leaves, there is a skew towards increased TE expression. The largest TE up-regulation occurs in the tissues of the male reproductive lineage, including unicellular and bicellular microspores, mature pollen and isolated sperm cells (Fig 3A). The number of up-regulated TEs compared to down-regulated TEs in these tissues suggest that there is a genome-wide activation of TE expression, similar to the DRTS event that occurs in Arabidopsis pollen [31,36]. One important distinction is that TE expression is present in maize sperm cells (Fig 3A), whereas it is not detected in Arabidopsis sperm cells [31]. To verify this finding, we compared our sperm cell RNA-seq data to an independent maize sperm cell dataset [30]. We found that TEs are also significantly expressed in this independent dataset, and 70% of those expressed TEs are also detected in our dataset (p<0.001) (Fig 3B). This shared set of 810 sperm cell-expressed TEs (38% of those detected in our dataset), supports the conclusion that significant expression of TEs occurs in maize sperm cells. Of the sperm cell-expressed TEs, 36% were not observable in total pollen, but rather required the isolation and enrichment of sperm cells for detection (Fig 3B). Overall, we detect 157 TEs expressed in both sperm cell datasets that are not expressed throughout development, but specifically in the sperm cells (sperm-cell exclusive).

**Fig 3.**
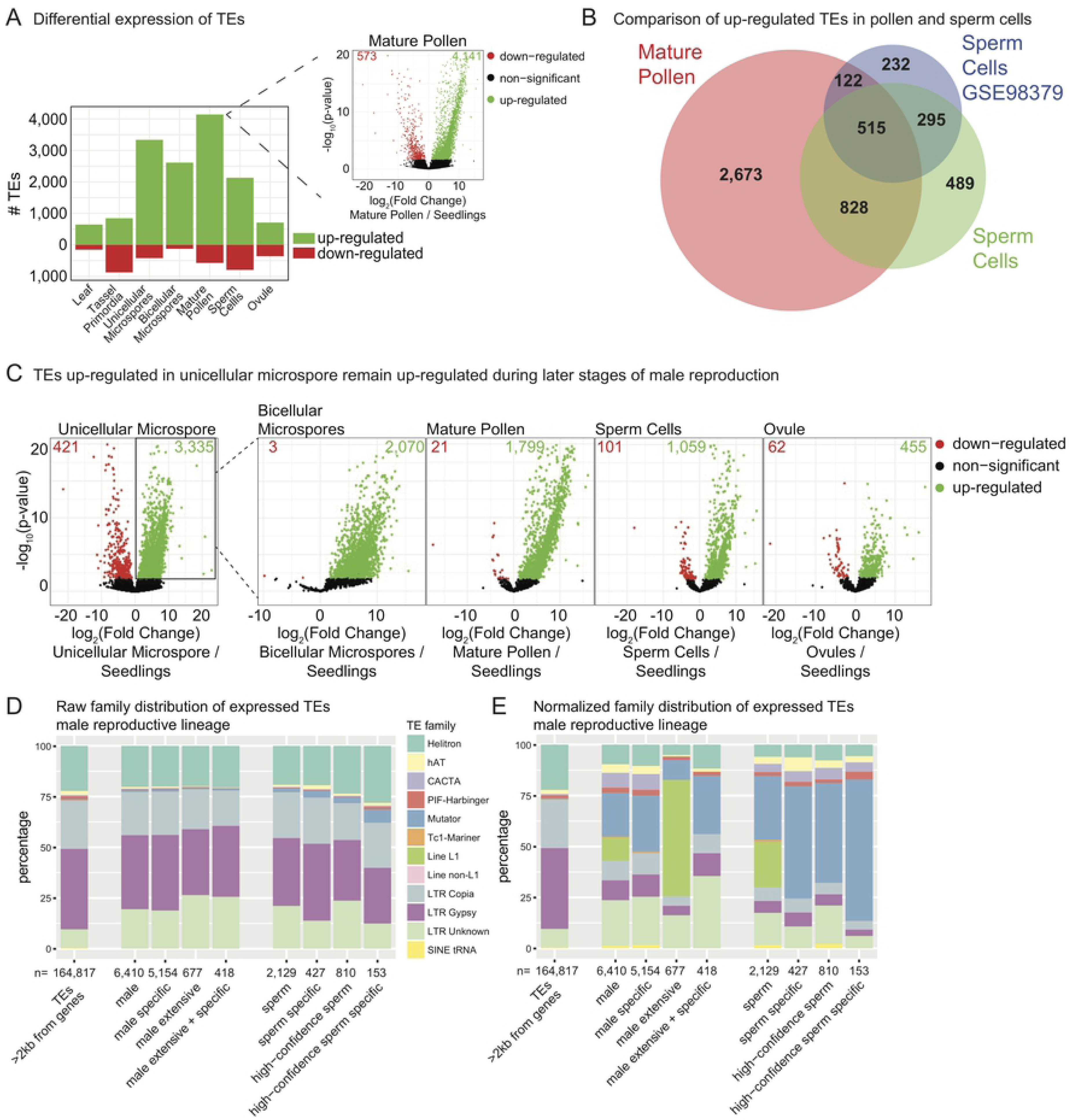
High TE expression in the maize male gametophyte lineage. **(A)** Number of differentially expressed TEs in seven tissues compared to seedlings. The inset volcano plot shows for mature pollen how differentially expressed TEs were identified. Green and red numbers within the volcano plot indicate how many TEs were statistically up- or down-regulated, respectively. **(B)** Number of up-regulated TEs in mature pollen compared to isolated sperm cells from this study and a previously published distinct isolation and sequencing of sperm cell mRNA. **(C)** Starting with TEs differentially up-regulated in unicellular microspores (boxed, far left volcano plot), we determined how many of these same TEs are expressed at other developmental time points. **(D)** Raw distribution of expressed TE family annotations. ‘Male’ refers to the set of TEs expressed in any male lineage dataset (MS, MS-B, MP, SC). ‘Male specific’ are TEs expressed in only the male lineage (not other tissues / timepoints). ‘Male extensive’ TEs are expressed in all of the male lineage datasets, and ‘male extensive + specific’ refers to TEs expressed all male lineage datasets and not other tissues / timepoints. ‘High-confidence sperm’ refers to TEs identified as expressed in both analyzed sperm cell datasets from part B. **(E)** Expressed TE family annotations normalized to the genome-wide TE distribution of TEs >2 kb from genes. Categories are the same as part D.

A second notable difference between maize and Arabidopsis is the activation of TE expression early in the male gametophytic phase of maize. A genome-wide increase in TE transcript levels is detected at the earliest post-meiotic stage tested, the microspore, in contrast to low TE expression in the sporophytic tassel primordia (Fig 3A). Arabidopsis TE expression occurs only late in pollen development, after pollen mitosis I when the somatic vegetative cell is generated [31]. To determine if TEs were indeed activated early in maize male reproductive development, we asked if the same TEs that we identified as expressed in the unicellular microspore remain active throughout the male reproductive lineage. We used the set of differentially expressed up-regulated TEs in unicellular microspores (3,335) and found that 62% are still expressed in bicellular microspores and 54% in mature pollen (Fig 3C), demonstrating that once TEs are activated early in development, expression and/or steady-state mRNA frequently remains through pollen maturation. Only some of these male-lineage expressed TEs continue to be expressed in sperm cells (32%), raising the possibility that many TEs with active expression in the early gametophytic stages are under negative/repressive regulation in the gametes. This large-scale developmental activation is potentially limited to the male lineage, as ovules express relatively few TEs (Fig 3A) and only 14% of the male lineage-expressed elements (Fig 3C). Together, our data demonstrate conserved activation of TE expression in the male gametophytes of maize and Arabidopsis, with key differences such as the developmental timing and localization of TE expression in the gamete cells.

We determined what types of TEs activate in the male reproductive lineage and sperm cells and compared these to the whole-genome distribution of TEs analyzed. Overall, both male lineage-expressed TEs and sperm cell-expressed TEs reflect the genome-wide TE distribution (Fig 3D). This suggests that TE family type does not have a determining role in the developmental regulation of TE expression. One notable exception is the enrichment of *Mutator* family TE expression in sperm cells (Fig 3D). When normalized for genome-wide TE distribution, *Mutator* element expression is highly enriched across the male lineage, including in sperm cells (Fig 3E). The expression of some *Mutator* TEs in sperm cells is both high confidence (present in both sperm cell datasets) and specific to only that tissue (high confidence sperm cell specific, Fig 3E). LINE L1 elements are also expressed throughout the male lineage and sperm cells, but their expression is general and not specific to these cell types (Fig 3E). Our data demonstrate that there is a general (TE family-independent) activation of TE expression in the male reproductive lineage, with one observable bias towards *Mutator* family expression in both the male lineage and sperm cells.

### Mature pollen and sperm cells display coexpression of highly expressed genes and their neighboring TEs

To determine if TEs have an effect on neighboring gene expression, or vise versa, we analyzed the 36,945 assayable TEs within 2kb of genes from Fig 2A. We calculated the absolute expression level of each genic isoform and categorized them into 100 bins of expression levels for each developmental stage (Fig 4). We find no relationship between gene expression level and the number of up- or down-regulated TEs in tassel primordia or microspores (top row, Fig 4A). In contrast, in both mature pollen and isolated sperm cells there is a positive association between highly expressed genes and the number of up-regulated TEs within 2kb of those genes (bottom row, Fig 4A). Similarly, there is a negative correlation between high gene expression and the number of down-regulated TEs in pollen and sperm cells (Fig 4A). This relationship is not due to the fact that pollen or sperm cell-expressed genes are more likely to be located nearby a TE (S3A Fig). We confirmed that this association between TE and gene regulation in sperm and mature pollen is not due to sample contamination between these two datasets (S3B Fig). It is unclear from these data whether gene expression is influencing TE expression, or TE expression is affecting gene regulation. However, we conclude that specifically in the mature male gametophyte the highest expressed genes are near actively expressing TEs.

**Fig 4.**
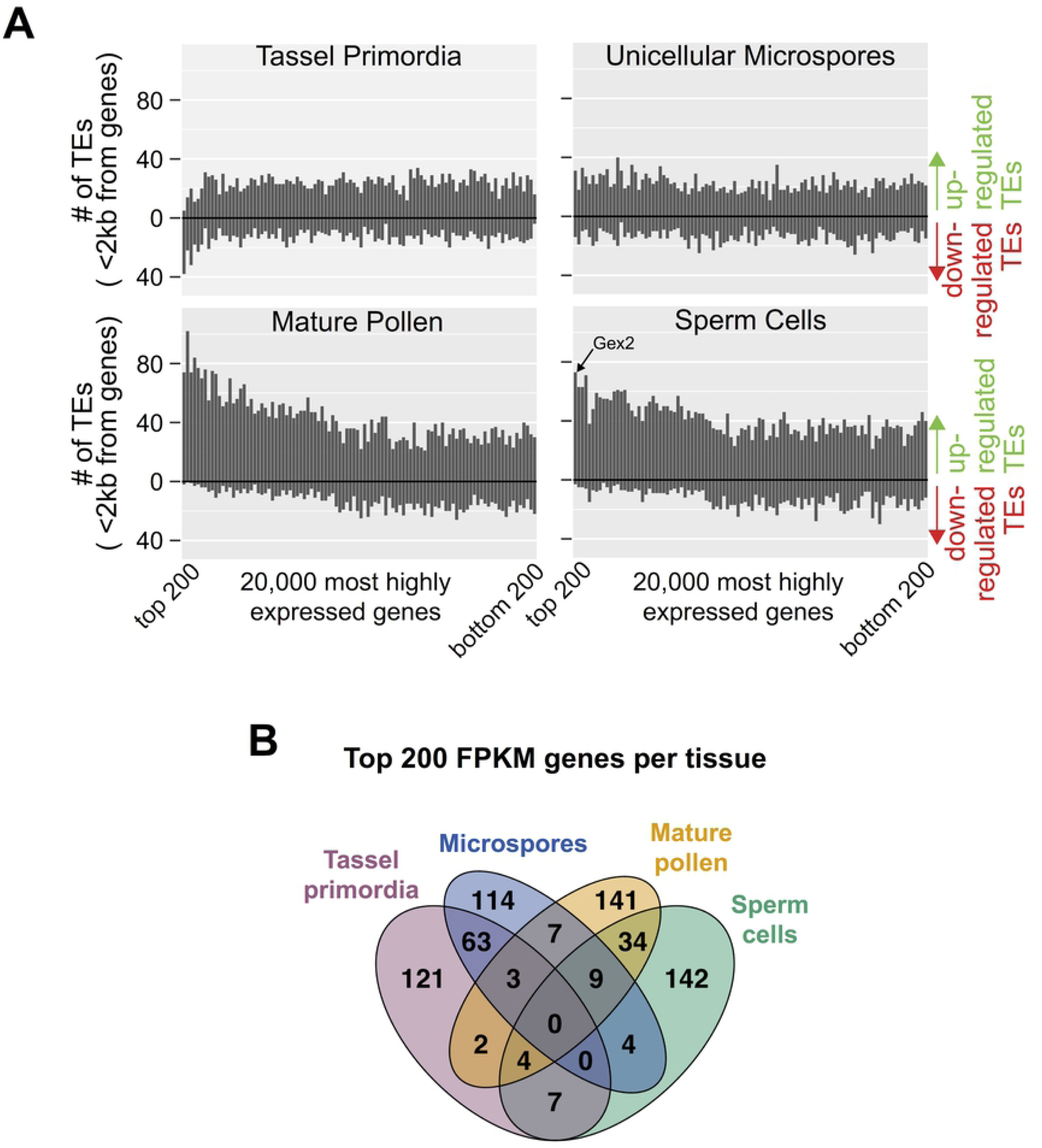
Co-regulation of TE and gene expression in the male gametophyte. **(A)** For each tissue type, the top 20,000 most highly expressed genes are distributed along the X-axis in bins of 200, with the most highly expressed bin on the far left. For each bin the number of up- and down-regulated TEs near (<2kb) its genes is then summed on the Y-axis. In unicellular microspores (top right) there is little correlation, while in mature pollen (bottom left) the most highly expressed genes are near primarily up-regulated TEs. The bin location of *gex2* (see Fig. 7) is annotated in the sperm cell data (bottom right). **(B)** High expression levels are associated with developmental specificity: approximately 2/3 of the genes associated with the highest FPKM values in each of the four sample types are highly expressed in only that sample type.

Comparison of the most highly expressed genes from mature pollen and sperm cells with other tissues assessed in this study (tassel primordia and microspores) shows that such transcripts were generally associated with high tissue-specificity (Fig 4B). Among the top 200 highly expressed genes by FPKM value, two-thirds of the genes in each tissue were highly expressed only in that tissue. No single gene was highly expressed in all four tissues. These data are consistent with the idea that genes highly expressed at a particular developmental stage contribute a genetic function specifically required at that stage. We next aimed to determine if these highly-expressed genes, many of which are adjacent to expressed TEs, have measurable functional roles in the male gametophyte.

### Large-scale insertional mutagenesis supports a relationship between transcript level and fitness contribution for vegetative cell-expressed genes

The developmental gene expression dataset generated by this study provided a quantitative framework in which to assess gene function, testing the hypothesis that highly expressed genes contribute significantly to reproductive success – i.e, fitness. The functional validation approach we used relied on a large, sequence-indexed collection of green fluorescent protein (GFP)-marked transposable element (*Ds-GFP*) insertion mutants [44], enabling assessment of the effects of mutations in select genes (Fig 5). We focused on expression data from the MP and SC stages, as these display coordinated expression with TEs (Fig 4). In addition, these stages have distinctive cell fates and roles in reproduction: the vegetative cell generates the pollen tube for competitive delivery of gametes, and the sperm cells accomplish double fertilization. Expression data from seedlings [26] was used to design a comparator sporophytic control. Highly expressed genes, operationally defined as in the top 20% for a tissue by FPKM, were grouped into three mutually exclusive classes: Seedling, Sperm Cell, and Vegetative Cell. The Seedling group also excluded any gene highly expressed in either MP or SC. Due to the significant overlap among genes highly expressed in both MP and SC, we compared expression values to assign each of these genes to a single class. Vegetative Cell genes were not only highly expressed in MP, but were also associated with an FPKM greater in MP than in SC, and vice versa for Sperm Cell genes (S5 Table). All genes in these classes were then cross-referenced with *Ds-GFP* insertion locations to identify potential mutant alleles for study, restricting the search to insertions in coding sequence (CDS), as these were rationalized as most likely to generate loss-of-function effects. Finally, to insure our results were as generalizable as possible, each class list was randomized to identify the specific subset of *Ds-GFP* lines for study. Insertion locations were verified by PCR for 64 of 83 alleles obtained (S6 Table) (see Methods), of which 56, representing mutations in 52 genes, generated sufficient transmission data to include in our final analysis.

**Fig 5.**
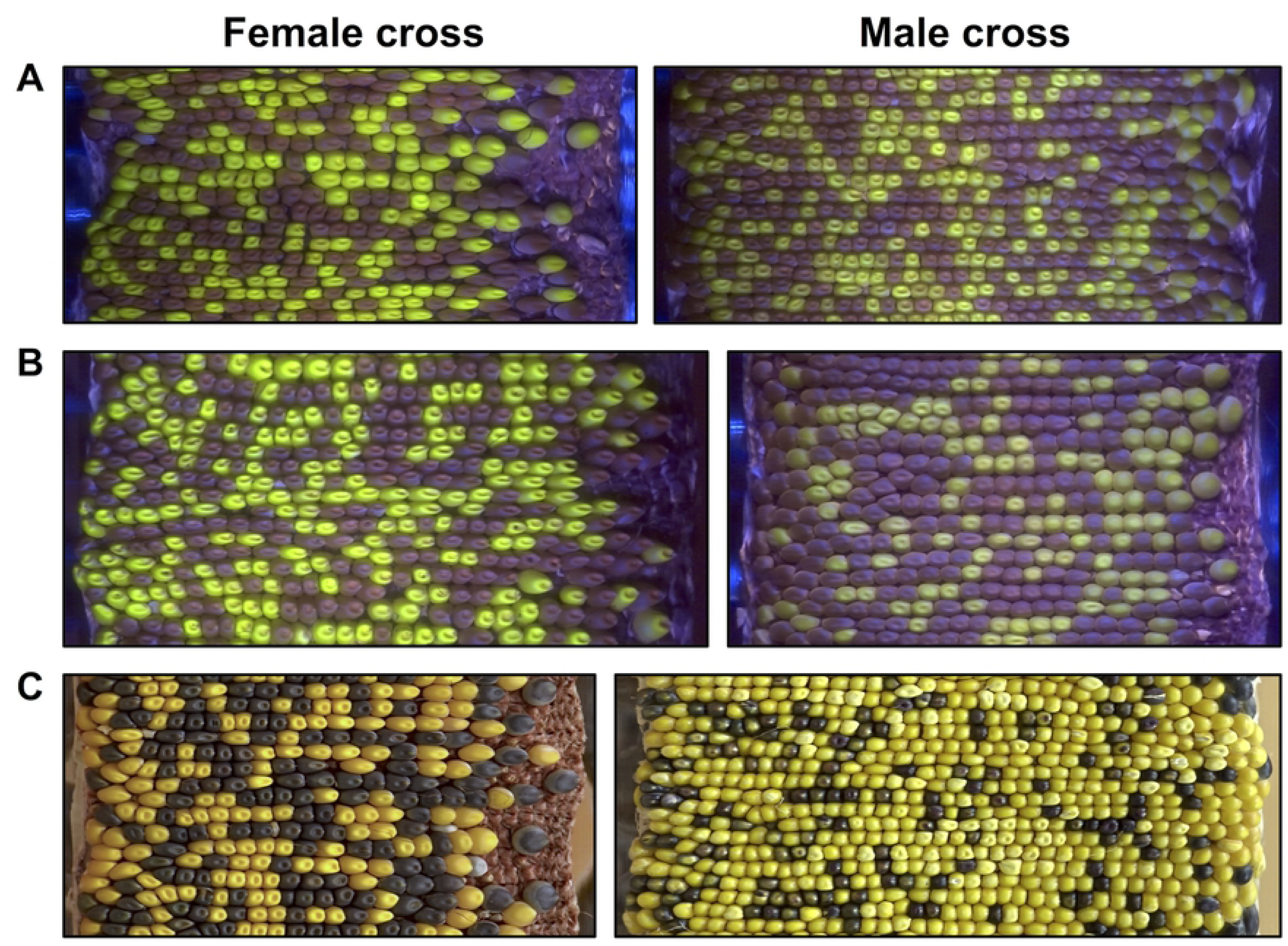
Large-scale tracking of seed marker transmission frequencies was accomplished by generating ear projections with a custom built rotational scanner. When crossed either through the male or the female, *Ds-GFP* mutant allele *tdsgR107C12* (in gene Zm00001d012382), marked by green fluorescent seeds, shows 1:1 Mendelian inheritance (50% transmission of the GFP seed marker). Images captured in blue light with an orange filter. Mutant alleles in other genes, such as *tdsgR102H01* (Zm00001d037695), showed non-Mendelian segregation when crossed through the male (37.5% GFP transmission). Segregation through the female remained Mendelian, indicating a male-specific transmission defect. **(C)** For some mutant alleles (∼10% of lines in this study), the anthocyanin transgene *C1* was tightly linked to the insertion mutant. In these cases, seeds carrying a mutant allele of a gene of interest could be tracked by their purple color. Here, insertion *tdsgR96C12* (Zm00001d015901) shows a strong male-specific transmission defect (24.8% *C1* transmission through the male). Images captured in full spectrum visible light.

Mendelian inheritance predicts 50% transmission of mutant and wild-type alleles when a heterozygous mutant is outcrossed to a wild-type plant. However, a mutation that alters the function of a gene expressed during the haploid gametophytic phase can result in a reduced transmission rate if that gene contributes to the fitness of the male gametophyte – i.e., to its ability to succeed in the highly competitive process of pollen tube growth, given that 50% of the pollen population will be wild-type for the same gene. Thus, reduced transmission of a mutant through the male (a male transmission defect) provides not only evidence for gene function in the gametophyte, but also a measure of the mutated gene’s contribution to fitness. Transmission rates through the female serve as a control, as 50% transmission through the female would confirm both a single *Ds-GFP* insertion in the genome and male-specificity for any defect identified. To measure the fitness cost of each *Ds-GFP* insertion, heterozygous mutant plants were reciprocally outcrossed with a heavy pollen load to a wild-type plant, maximizing pollen competition within each silk. Transmission rates were then quantified by assessing the ratio of the non-mutant to mutant progeny using a novel scanning system and image analysis pipeline (Fig 5) (see Methods) [45]. Mutant alleles were tracked using linked endosperm markers: either the GFP encoded by the inserted transposable element (Fig 5A-B), or, in ∼10% of the lines, a tightly linked *C1*^*+*^ anthocyanin transgene (present due to the initial *Ds-GFP* generation protocol) (Fig 5C, S7 Table). For an allele to be included in the final dataset, we required a minimum of three independent male outcrosses from two different plants. The number of seeds categorized for each allele ranged from 1,522 to 5,219, with an average of 2,807.

Transmission rates for all groups were tested through quasi-likelihood tests on generalized linear models with a logit link function for binomial counts (see Methods, S8 Table). When crossed through the female, no genes showed significant differences from Mendelian inheritance (Fig 6A). When crossed through the male, no genes with insertion alleles in the Seedling category (n=10) showed evidence of abnormal transmission rates (Fig 6B). Most Sperm Cell genes (n=10, 90%) showed no statistically significant transmission defects, with one notable exception (two independent alleles of the *gex2* gene, described in detail below) (Fig 6C). However, among Vegetative Cell genes tested (n=32), a larger proportion of insertion alleles (7 out of 32 or 21.9%) showed significant male transmission defects (quasi-likelihood test, adjusted p-value threshold < 0.05) (Fig 6D). The proportions of genes with transmission defects in the three classes were not significantly different by Fisher’s exact test (Seedling vs Sperm Cell p-value = 0.500, Seedling vs Vegetative Cell p-value = 0.125, Vegetative Cell vs Sperm Cell p-value = 0.374), likely due to the small number of mutations assessed in the Seedling and Sperm Cell classes.

**Fig 6.**
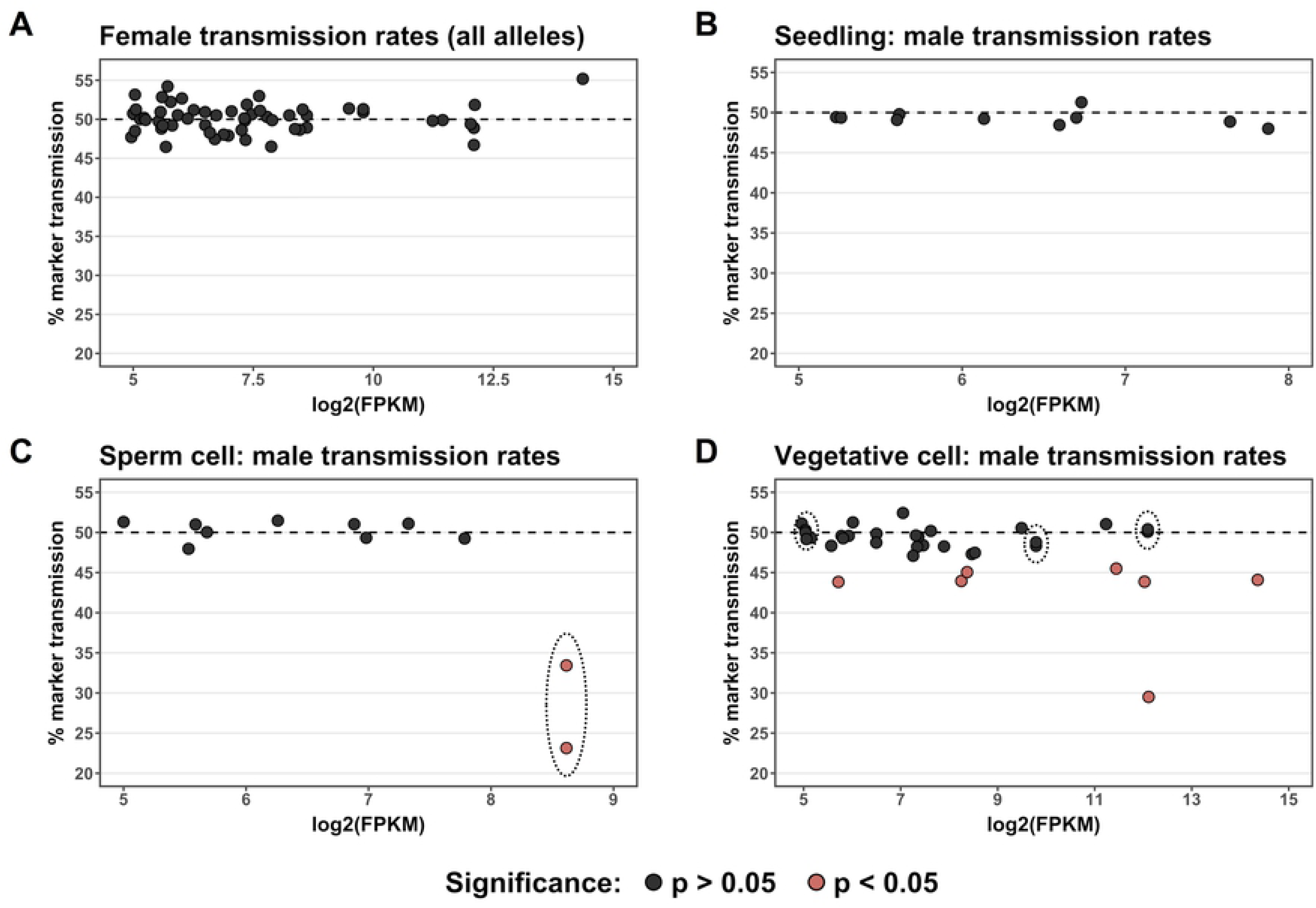
Highly expressed pollen genes are more likely to be associated with decreased male transmission rates. Alleles with CDS insertions were tested for differences from Mendelian inheritance using a quasi-likelihood test, with p-values corrected for multiple testing using the Benjamini-Hochberg procedure; alleles with quasi-likelihood adjusted p-value < 0.05 are represented in pink. Alleles are plotted by the log_2_(FPKM) of their respective gene according to that gene’s expression class (Seedling, Vegetative Cell, or Sperm Cell). Insertion alleles distributed among the classes as follows: Vegetative Cell, 35 alleles; Sperm Cell, 11 alleles; Seedling, 10 alleles. Genes represented by two independent insertion alleles are enclosed by dotted lines. **(A)** Transmission rates of 56 mutant allele seed markers for heterozygous *Ds-GFP* mutant plants crossed through the female. **(B)** Transmission rates for alleles in the negative control Seedling class when crossed through the male. **(C)** For genes belonging to the Sperm Cell class, one out of ten (10%) was associated with significant non-Mendelian inheritance. The single gene with a male transmission defect in this group (*gex2*) showed a strong defect for both of the independent alleles tested. **(D)** An increased proportion of the genes in the Vegetative Cell class were associated with significant non-Mendelian inheritance when mutant (7/32 genes, 21.9%). In this class, an increase in log_2_(FPKM) was significantly associated with a decrease in marker transmission (linear regression, p = 0.0120).

The majority of transmission defects in the Vegetative Cell class genes (six of the seven with significant effects) were modest, at approximately 45% transmission, with only one reducing transmission strongly, to ∼30%. Two of the seven Vegetative Cell genes with transmission defects were adjacent to TEs (S7 Table). Notably, all but one of the genes associated with significant defects were measured at a log_2_(FPKM) > 8 (i.e., in the top 5% of Vegetative Cell genes by FPKM). Above this expression level, the percentage of genes associated with a significant defect rises to 50% (six out of twelve). Thus, the most highly expressed Vegetative Cell genes are significantly more likely to be associated with non-Mendelian transmission than the group of Vegetative Cell genes below this expression threshold (1 out of 20) (Fisher’s exact test, p-value = 0.00572). Consistent with this observation, an increase in log_2_(FPKM) was associated with both reduced transmission rate and an increase in -log_10_(p-value) (linear regression, p-value = 0.0120, 0.0255, respectively; adjusted R^2^ = 0.151, 0.116, respectively). Thus, our data suggest that higher transcript level in the Vegetative Cell predicts a gene-specific contribution to male gametophytic fitness. Vegetative Cell genes showing non-Mendelian inheritance had a range of predicted cellular functions, including cell wall modification, cell signaling, protein folding, vesicle trafficking, and actin binding (Table 1).

**Table 1.**
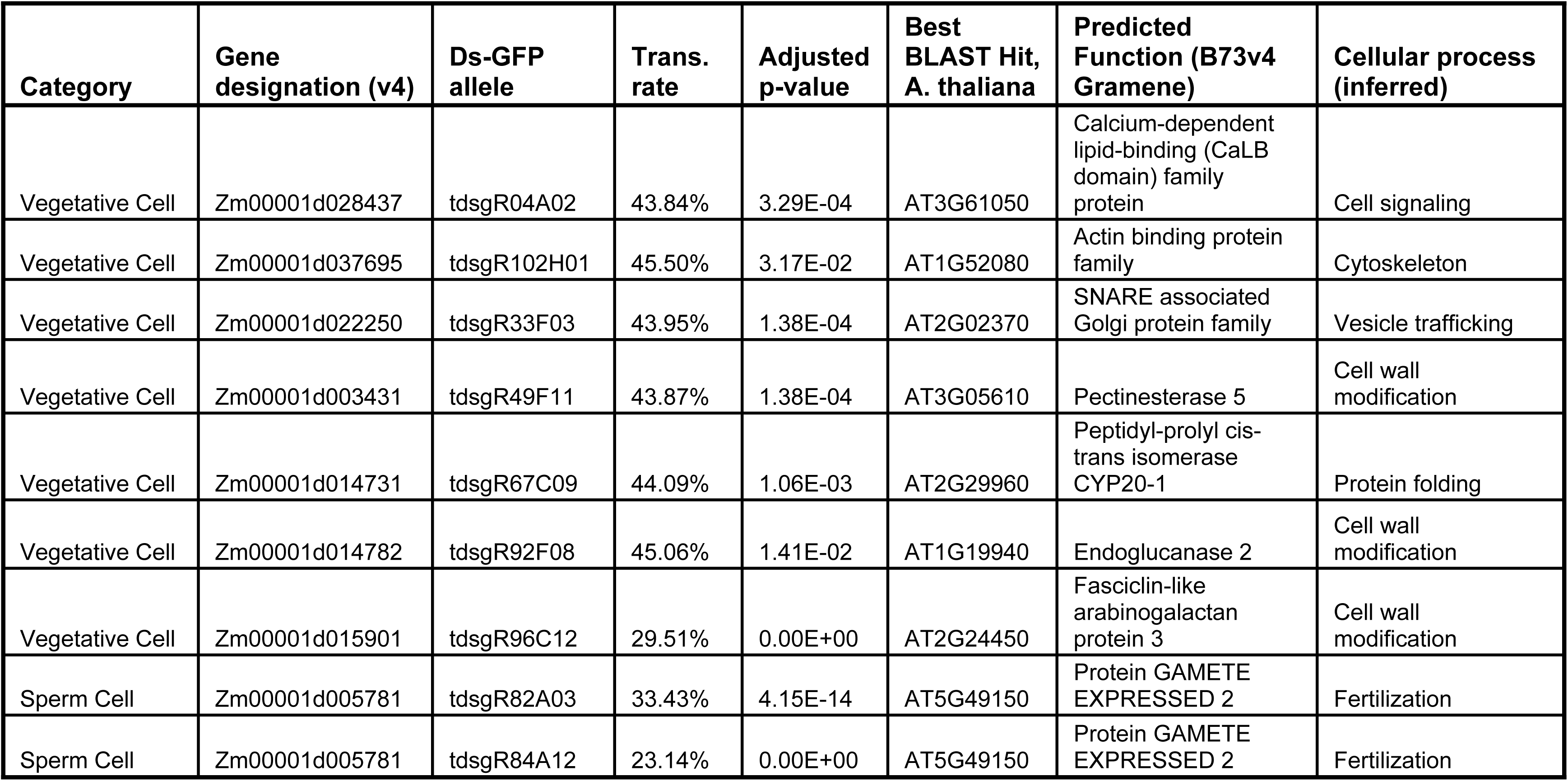
Characteristics of genes showing non-Mendelian inheritance.

| Category | Gene designation (v4) | Ds-GFP allele | Trans. rate | Adjusted p-value | Best BLAST Hit, *A. thaliana* | Predicted Function (B73v4 Gramene) | Cellular process (inferred) |
| --- | --- | --- | --- | --- | --- | --- | --- |
| Vegetative Cell | Zm00001d028437 | tdsgR04A02 | 43.84% | 3.29E-04 | AT3G61050 | Calcium-dependent lipid-binding (CaLB domain) family protein | Cell signaling |
| Vegetative Cell | Zm00001d037695 | tdsgR102H01 | 45.50% | 3.17E-02 | AT1G52080 | Actin binding protein family | Cytoskeleton |
| Vegetative Cell | Zm00001d022250 | tdsgR33F03 | 43.95% | 1.38E-04 | AT2G02370 | SNARE associated Golgi protein family | Vesicle trafficking |
| Vegetative Cell | Zm00001d003431 | tdsgR49F11 | 43.87% | 1.38E-04 | AT3G05610 | Pectinesterase 5 | Cell wall modification |
| Vegetative Cell | Zm00001d014731 | tdsgR67C09 | 44.09% | 1.06E-03 | AT2G29960 | Peptidyl-prolyl cis-trans isomerase CYP20-1 | Protein folding |
| Vegetative Cell | Zm00001d014782 | tdsgR92F08 | 45.06% | 1.41E-02 | AT1G19940 | Endoglucanase 2 | Cell wall modification |
| Vegetative Cell | Zm00001d015901 | tdsgR96C12 | 29.51% | 0.00E+00 | AT2G24450 | Fasciclin-like arabinogalactan protein 3 | Cell wall modification |
| Sperm Cell | Zm00001d005781 | tdsgR82A03 | 33.43% | 4.15E-14 | AT5G49150 | Protein GAMETE EXPRESSED 2 | Fertilization |
| Sperm Cell | Zm00001d005781 | tdsgR84A12 | 23.14% | 0.00E+00 | AT5G49150 | Protein GAMETE EXPRESSED 2 | Fertilization |

To ensure the experimental design was robust, we examined two potential confounding variables: the presence of the *wx1-m7::Ac* allele in a subset of lines tested and the potential for epigenetic silencing of GFP transgenes (see S1 Methods). We found no evidence that the presence of *wx1-m7::Ac* significantly impacted the overall conclusions drawn from the dataset, as well as no evidence of epigenetic silencing of GFP transgenes.

### Insertional mutants in sperm cell-expressed *gex2* cause paternally triggered aberrant seed development

The male-specific transmission defect for the sole affected gene in the Sperm Cell class, Zm00001d005781 (GRMZM2G036832), was significantly higher than the average defect across all genes identified with decreased transmission through the male. Sequencing confirmed that the *Ds-GFP* elements in the two independent alleles, tdsgR82A03 and tdsgR84A12, associated with average transmission rates of 33.4% and 23.1%, respectively, were inserted into their predicted exonic locations (Fig 7A). In addition, both alleles showed an unusual phenotype of underdeveloped or aborted seeds and ovules with no apparent seed development when crossed through the male, despite heavy pollination (Fig 7B). These features motivated further investigation of this gene.

**Fig 7.**
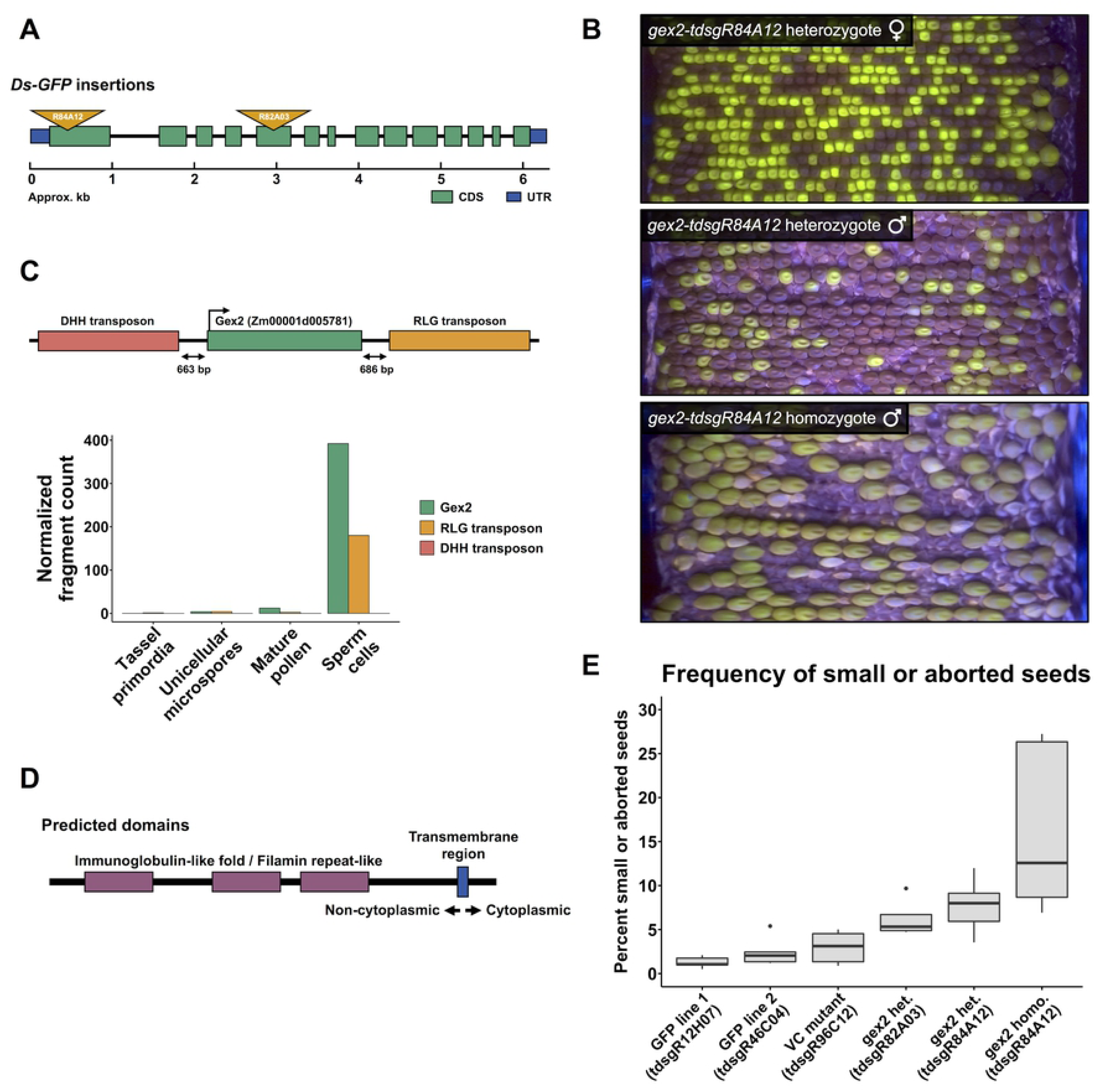
Mutations in the sperm cell-specific *gex2* gene cause aberrant seed development. **(A)** The exon/intron structure of *gex2* (Zm00001d005781/GRMZM2G036832), showing the locations of the two independent *Ds-GFP* insertion mutants. **(B)** Ear projections of *gex2* mutant outcrosses. Top: heterozygote outcrossed as female, showing 1:1 transmission of the GFP-tagged allele. Middle: heterozygote outcrossed as a male, with 26.1% transmission of the mutant allele. Additionally, small seeds and occasional, small gaps between seeds are visible. Bottom: homozygous mutant outcrossed as a male, with many small seeds and large gaps, despite heavy pollination. **(C)** Genomic neighborhood of the GEX2 locus, with two nearby TEs, and their RNA-seq expression levels across male reproductive development. **(D)** Predicted domain structure of GEX2. **(E)** Quantification of small/aborted seeds resulting from pollination by *gex2* mutant plants and controls. Controls included two *Ds-GFP* lines that did not show transmission defects (*tdsgR12H07* and *tdsgR46C04*) and one *Ds-GFP* line that showed a strong transmission defect in the vegetative cell group (*tdsgR96C12*). A higher percentage of small/aborted seeds was present following pollination by heterozygous *gex2* plants representing the two mutant alleles (*tdsgR82A03* and *tdsgR84A12*), and pollination by homozygous *gex2-tdsgR84A12* plants further increased the percentage of small/aborted seeds.

Across maize tissues, Zm00001d005781 is highly and specifically expressed in sperm cells [41] (Fig 7C). Zm00001d005781 was previously identified in maize sperm cells via EST sequencing and named *gamete expressed 2*, or *gex2*, but no mutant has been described [46]. The gene is among the top 200 expressed in sperm cells, and like many highly tissue-specific genes expressed in pollen and sperm, it is within 2kb of a transcriptionally active TE, both a downstream RLG retrotransposon that displays sperm cell-specific activation and an upstream DHH family TE that is not detectably transcribed (Fig 7C). Predicted *gex2*-like genes are widely distributed throughout the currently sequenced Embryophyta taxa. The Arabidopsis ortholog, *GEX2*, has been described as necessary for gamete attachment and effective double fertilization [47]. Maize *gex2*, a single copy gene, encodes a protein with predicted domain structure similar to that of Arabidopsis GEX2 (44.2% protein similarity overall). Both proteins harbor a large N-terminal non-cytoplasmic region including filimin repeat-like domains, predicted transmembrane domains near their C-termini, and a small C-terminal cytoplasmic region (Fig 7D).

Small and aborted seeds were quantified for both *gex2* insertion alleles when outcrossed to wild-type plants as heterozygotes, as well as for outcrosses from *gex2-tdsgR84A12* homozygotes (S10 Table, S4 Fig). As controls, the same assessment was made for pollinations from two different heterozygous *Ds-GFP* insertion lines that were not associated with transmission defects (*tdsgR12H07, tdsgR46C04*), as well as heterozygous plants carrying the *Ds-GFP* associated with the strongest male transmission defect (29.5% transmission) in the Vegetative Cell class (*tdsgR96C12*). These three *Ds-GFP* insertions showed similar percentages of aborted seeds (Fig 7E). In contrast, pollination with both *gex2::Ds-GFP* insertion alleles was associated with increased percentages of small or aborted seeds, significantly so in *gex2-tdsgR84A12* (pairwise t-test against *Ds-GFP* controls separately, all p-values < 0.05). Pollination from *gex2-tdsgR84A12* homozygotes approximately doubled the percentage of small or aborted seed percentages from *gex2::Ds-GFP* heterozygous plants (pairwise t-test against *Ds-GFP* controls separately, all p-values < 0.01). From the heterozygous *gex2::Ds-GFP* crosses, small seeds with endosperm large enough for DNA preparation were genotyped, and 79.2% were found to harbor the *gex2* mutation, whereas in crosses from the *tdsgR46C04* control, the *Ds-GFP* insertion showed Mendelian segregation in small seeds. These data support the hypothesis that aberrant seed development is associated with fertilization by *gex2::Ds-GFP* sperm.

If GEX2 acts to promote double fertilization, the arrival of a *gex2::Ds-GFP* pollen tube at the embryo sac could lead to failure of one or both fertilization events. Given an active polytubey block, this could produce the observed gaps between seeds on the ear, resulting from ovules associated with completely failed fertilization, or with very early seed abortion due to single fertilization. To explore this possibility, seedless area was measured in ear projection images for ears pollinated by *gex2::Ds-GFP* heterozygous and homozygous mutants, as well as for the *Ds-GFP* control ears already described. As heterozygotes, all three *Ds-GFP* controls as well as the *gex2-tdsgR82A03* allele are associated with low levels of seedless area (each at 4%), whereas the *gex2-tdsgR84A12* shows a non-significant increase in seedless area (8.70%) (S5 Fig). However, ears pollinated by homozygous *gex2*-*tdsgR84A12* pollen had 31.48% seedless area, a significant increase (pairwise t-test against *Ds-GFP* controls separately, all p-values < 0.0001). To test for aberrant fertilization more directly, seed development was assessed at 4 days post-pollination with either wild-type or *gex2*-*tdsgR84A12* homozygous pollen (Fig 8, Table 2). Typical embryo and endosperm development, as well as indication of the polytubey block (i.e., arrival of only single pollen tubes at the embryo sac), was observed in all ovules assessed from wild-type pollination. In contrast, half of the ovules assessed following pollination with *gex2::Ds-GFP* showed significant evidence of abnormal double fertilization, demonstrating single fertilization of either embryo or endosperm or indication of arrival of more than one pollen tube at the embryo sac (Fisher’s exact test, p-value = 0.000241). We conclude that maize GEX2 is part of the sperm cell machinery that helps insure proper double fertilization.

**Table 2.**
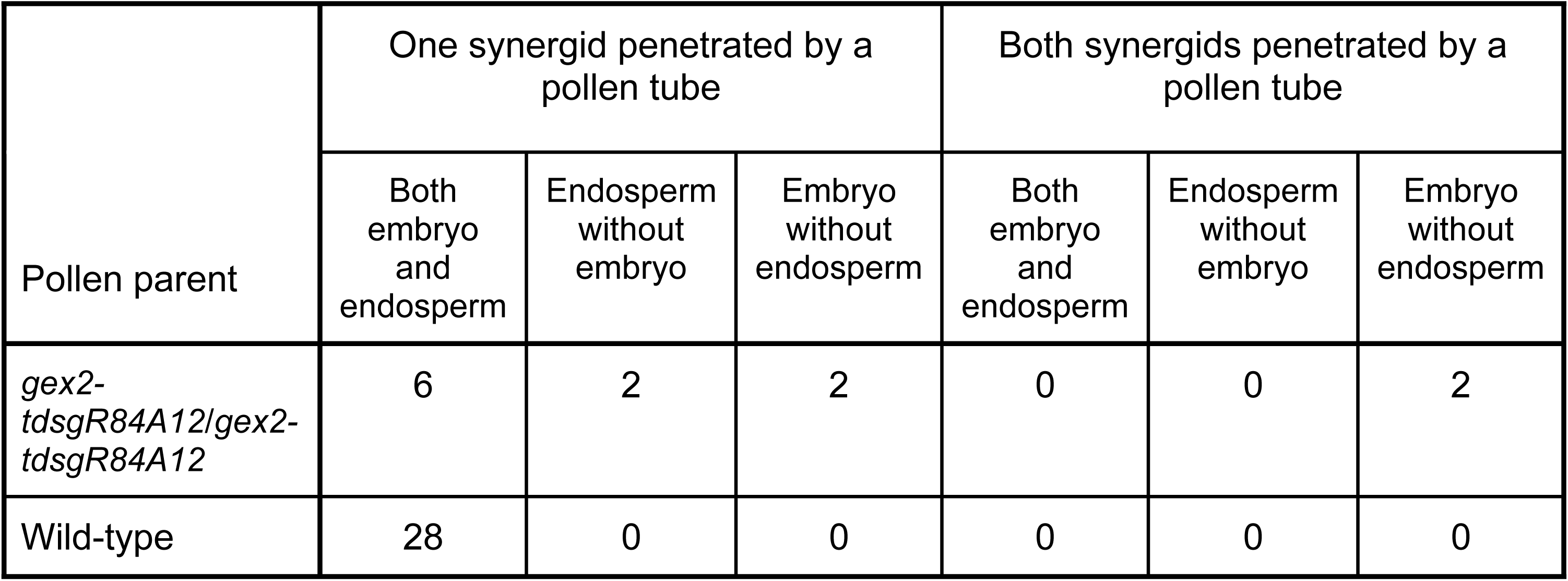
Seed development at 4 days after pollination by wild-type or gex2::Ds-GFP pollen.

| Pollen parent | One synergid penetrated by a pollen tube |  |  | Both synergids penetrated by a pollen tube |  |  |
| --- | --- | --- | --- | --- | --- | --- |
|  | Both embryo and endosperm | Endosperm without embryo | Embryo without endosperm | Both embryo and endosperm | Endosperm without embryo | Embryo without endosperm |
| *gex2-tdsgR84A12/gex2-tdsgR84A12* | 6 | 2 | 2 | 0 | 0 | 2 |
| Wild-type | 28 | 0 | 0 | 0 | 0 | 0 |

**Fig 8.**
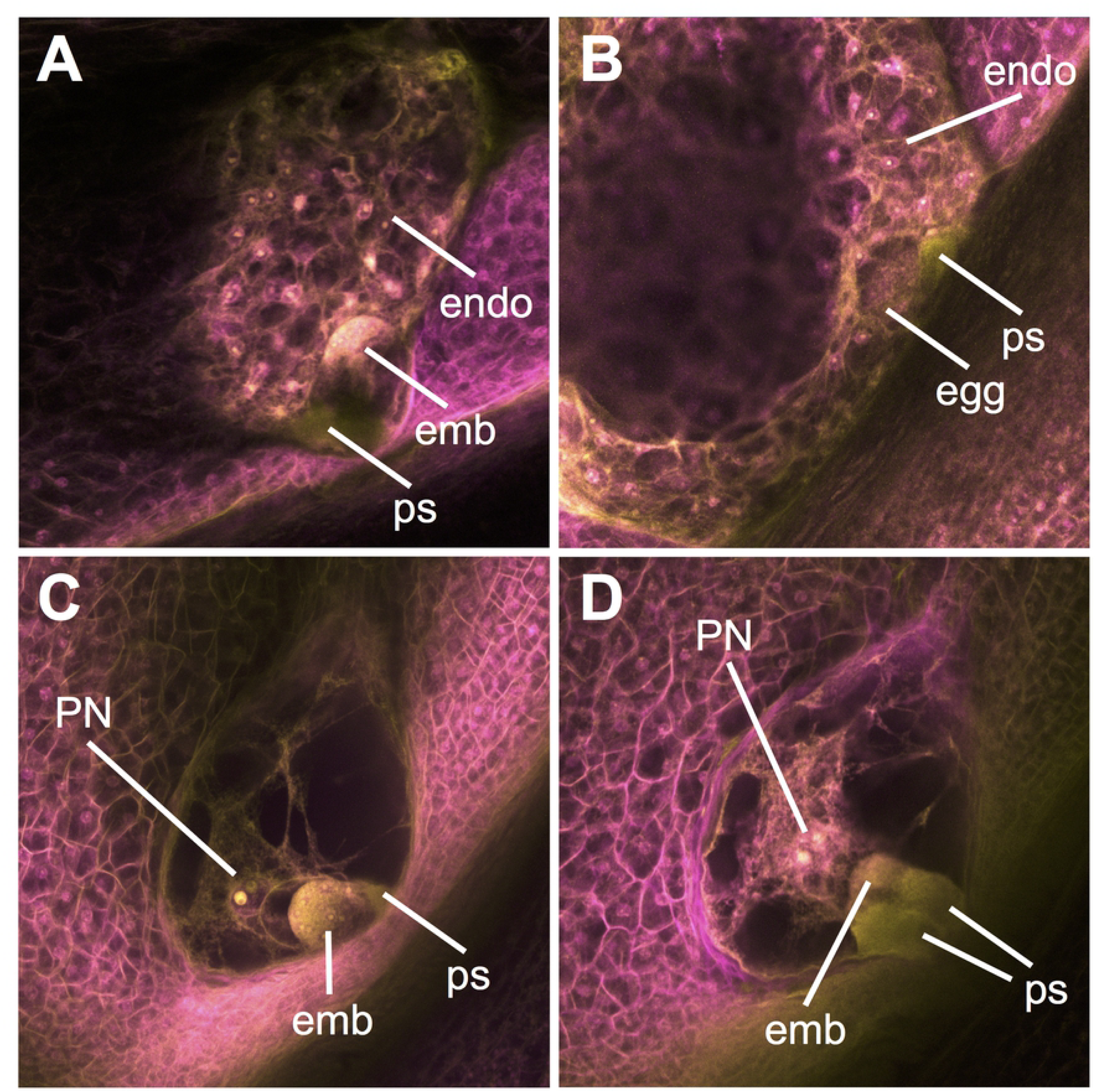
Pollination by *gex2-tdsgR84A12* leads to aberrant fertilization events and developing seed phenotypes. **(A)** Seed development in a typical ovule pollinated by wild-type pollen, with one synergid penetrated by a pollen tube, and both embryo and endosperm development initiated. **(B-D)** Abnormal phenotypes seen following *gex2* pollination. **(B)** Ovule with developing (cellularizing) endosperm but unfertilized egg cell. **(C)** Ovule with developing embryo but unfertilized central cell. **(D)** Ovule with both synergids penetrated by a pollen tube, and a developing embryo and unfertilized central cell. emb=embryo; endo=endosperm; ps=synergid penetrated by a pollen tube; PN=polar nuclei.

## Discussion

### Transposable element dynamics in the maize male gametophyte

Our analysis of TE expression during maize male reproductive development provides an informative comparison to similar analyses in Arabidopsis, an evolutionarily distant plant with a genome landscape that is quite distinct from maize. Although maize has a higher number and percentage of its genome occupied by TEs compared to Arabidopsis, we found that only a fraction of maize TEs are developmentally dynamic with regards to transcript accumulation. These ‘dynamic’ TEs tend to be longer elements than average, and are enriched for *Mutator* family DNA transposons and ‘unknown’ classification LTR retrotransposons. From this dynamic TE set, we were able to identify individual elements that are expressed in a number of specific tissues. However, more globally, there is a trend towards activation of TE transcription over the course of the development of the mature male gametophyte. This finding confirms that both monocots and eudicots have developmental activation of TE expression in pollen [31]. This conservation suggests that the roles of TE and TE-induced small RNAs during reproductive development may also be conserved between monocots and eudicots [38,48,49]. Consistent with our findings, a recent study found that spontaneous retrotransposon mutations are much more frequent through the male than the female in certain maize lines [34].

Although TE activation is conserved in maize and Arabidopsis pollen, we have identified key differences in the timing and location. Maize TE activation is detected earlier (in the unicellular microspore) compared to when it is thought to occur in Arabidopsis [31]. Transcripts from these early-activated TEs in the microspores typically remain detectable through pollen development and in the mature pollen grain, which may be due to continued expression or transcript stability. A second distinction is the location of activation, which in Arabidopsis is confined to the pollen vegetative cell nucleus [31,35], whereas in maize also occurs in sperm cells. *Mutator* family TE transcripts are overrepresented in the pool of sperm-cell TE transcripts, suggesting that this family of TEs may have evolved (or co-opted) specific regulatory mechanism(s) such as an enhancer element that confers expression in this cell type.

We also examined the correlation between gene and TE expression. We found that generally there is no association between highly expressed genes and TE activation. However, in mature pollen and sperm cells there is a positive correlation: the more highly expressed a gene is, the more likely it is to have expressed TEs within 2kb. This tissue-specific correlation is not due to there being more TEs near pollen- or sperm-expressed genes. This demonstrates a developmentally specific co-regulation of gene and TE expression. Several potential mechanisms account for this observation. First, the programmed activation of TE expression may influence chromatin, enhancer, or other regulatory function that influences the neighboring genes. Second, the genes and TEs may be directly controlled by the same mechanism of large-scale epigenetic activation. The same mechanisms that repress TE activation may be responsible for limiting the expression of some genes to a single specific tissue or developmental time point. Third, the gene activation may control the expression of the neighboring TE. Ongoing studies aim to decipher these possibilities.

### The *gex2* gene has a conserved role in promoting double fertilization

One highly and specifically expressed gene in maize sperm cells with a nearby activated TE is *gex2*. The gene *gex2* is located between two transposable elements, one of which (an RLG retrotransposon) was also specifically transcriptionally activated in sperm cells. Mutations in *gex2* led not only to severely reduced transmission through the male, but also, in contrast to other mutations analyzed in this study, to paternally triggered post-fertilization defects, such as an increase in underdeveloped and aborted seeds and an increase in the seedless surface of the ear. *gex2* was first identified in maize by sperm cell EST sequencing [46], which led to identification of the orthologous gene in Arabidopsis, *GEX2*, and its sperm cell-specific promoter [50]. In Arabidopsis, seed abortion and single fertilization events were observed at increased frequency in *gex2* mutant-pollinated plants, both of the egg cell (leading to seeds that contained only an embryo) and of the central cell (leading to seeds that contained only endosperm) [47]. Our results are similar, with maize *gex2* mutant pollen causing single fertilization events in embryo sacs, small and aborted seeds, and leading to aberrant early seed phenotypes consistent with single fertilization. Mechanistically, Arabidopsis GEX2 appears to contribute to gamete attachment through interactions between plasma membrane-localized GEX2 in the sperm cell and either the female egg or central cells. The two orthologues share a predicted domain structure, including a large N-terminal non-cytoplasmic region containing filimin repeat domains potentially acting in gamete attachment [47], raising the possibility that maize GEX2 acts similarly during double fertilization. However, we cannot rule out additional functions for *gex2* in the earlier phase of pollen tube growth.

Few genes with clear fertilization-associated functions have been identified in maize. Three mutations which cause paternally-triggered aberrant (‘rough’) endosperm development have been isolated, but their corresponding genes have not been molecularly identified [51]. Two genes that are more clearly linked to double fertilization, *MTL*/*NLD*/*ZmPLA1* [17–19] and *ZmDMP* [20], have been identified based on their contributions to pollen-induced haploid induction associated with the ‘Stock 6’ line of maize. In our RNA-seq dataset, *MTL*/*NLD*/*ZmPLA1* is detected at low levels in SC, but is 7-fold higher in MP, suggesting enriched expression in the vegetative cell; *ZmDMP* is not detected in either SC or MP. Similar to *gex2*, mutants in both *MTL*/*NLD*/*ZmPLA1* and *ZmDMP* cause small/aborted seed phenotypes when used as a pollen parent, raising the possibility that the three genes could act in related fertilization mechanisms. However, the exact mechanisms by which mutations in *MTL*/*NLD*/*ZmPLA1* and *ZmDMP* lead to haploid induction are unclear. Single fertilization of the central cell has been suggested as a possible mechanism [52], but more recent work points toward degradation of the paternal chromosomes in the zygote following double fertilization [53–55], while some data support the idea that both mechanisms contribute [56]. Some of the abnormal and early aborted seeds in *gex2* mutant pollinations are not explained simply by single fertilization events, which are expected to produce germless or endospermless seeds. The latter of these abort early and resemble unfertilized ovules [57], and thus would likely contribute to the observed gaps on the ears. Abnormal seeds with some endosperm development may indicate other post-fertilization defects caused by *gex2* loss-of-function, similar to *mtl/nld/zmpla1* or *zmdmp* loss-of-function. Direct, detailed comparison of seed phenotypes induced by pollination with these mutants could prove useful to understanding mechanisms both of pollen-induced haploid induction, and of fertilization in general.

### Maize pollen provides a powerful model for quantifying gene-specific contributions to fitness

Despite the explosion of omic-scale methods to characterize genomes and to measure molecular characters (e.g., transcript levels), our ability to predict phenotypic relevance for specific genes is limited, particularly in multicellular organisms with complex genomes. A simple assumption is that a high transcript level at a particular developmental stage implies an important function for the associated gene at that stage, thus pointing toward a potential phenotypic role. However, this has been difficult to test at larger scales, as generating gene-specific quantitative phenotypic data is laborious and standardization can be challenging. This study begins to address this assumption with an initial systematic assessment of the functional relevance of highly expressed genes in maize pollen, taking advantage of the ease of reciprocal outcross pollination in maize, the availability of a sufficient number of marked and likely null mutations, and the development of an imaging technique that enables sensitive quantitation.

The progamic (i.e., post-pollination) phase of male gametophyte development can be thought of as comprising two stages. In the first stage, pollen tube growth through the silk, male gametophytes compete to be the first to reach the female gametophyte. Each pollen grain is an independent multicellular organism, often genetically-distinct from others in the population due to meiotic recombination, with its sole purpose to deliver the sperm cells for double fertilization. Thus, in an outcrossing plant with an extensive stigma and style like maize, there is likely a heightened context for competition among individuals in pollen populations [10]. We reasoned that genes involved in this stage could be identified by measuring fitness costs for mutations in genes highly expressed in the vegetative cell, which is responsible for pollen tube growth. We found that CDS-insertion alleles for 7 out of 32 (21.9%) tested genes in this class are associated with mild to moderate male-specific transmission defects. These results support the idea that competitive pollen tube growth requires a broad array of genes, as genes associated with demonstrated fitness contributions were assigned to a variety of predicted cellular functions. However, it is worth noting that three of the seven are predicted to directly influence modification of the cell wall (Table 1). Extrapolating from this dataset to the entire genome gives an estimate of 600 vegetative cell genes in maize leading to non-Mendelian segregation when inactivated, at least under competitive conditions.

In the second stage, following the arrival of the pollen tube at the embryo sac, the pollen tube must properly penetrate the embryo sac and release the sperm cells it contains, which then must fuse with the egg cell and central cell for double fertilization. Due to the presence of the polytubey block, the second stage contrasts with the first in that competition is likely minimal. Upon delivery, sperm cells must perform the tasks of adhesion, communication, fusion with the egg cell and central cell, and karyogamy. However, due to the lack of competition, minor problems with these processes may not lead to failure. We aimed to investigate this second stage by testing genes highly expressed in the sperm cell. Our limited dataset identified 1 out of 10 of tested genes in this category as associated with a male transmission defect. This mutant gene, *gex2*, had among the strongest transmission defects observed in this study – and, intriguingly, is also associated with the highest FPKM among the sperm cell genes tested. Overall, our results suggest a scenario in which mutations in a larger proportion of genes operating during pollen tube growth lead to slight reductions in fitness, whereas mutations of relatively fewer genes operating in double fertilization lead to strong reductions in fitness. Testing a larger number of *Ds-GFP* insertions in vegetative cell- and sperm cell-enriched genes, encompassing a wider range of expression levels, would help establish if these trends are borne out.

To our knowledge, this study is the largest yet in plants to explore the possibility of a relationship between transcriptomic data and quantitative phenotypic effects of mutations. The largest phenomic studies (hundreds of mutant lines assessed) of leaf or reproduction-related characters using the Arabidopsis T-DNA mutant collection have identified phenotypic effects in ∼4% of lines assessed, although these were not guided by transcriptomic data [58,59]. Thus, the higher frequency of phenotypic effects we found could be due to sampling only mutations in genes that are most highly expressed at developmental stages most relevant to the phenotype. Genome scale measurement of the fitness costs of gene knockouts via competitive assays have been carried out in yeast [60][61] and bacteria [62][63]. All of these studies found that, for particular environmental conditions, there was little to no correlation between the expression level of a gene and its impact on fitness in that condition. In contrast, we found a small but significant correlation between high expression and fitness in our smaller dataset of vegetative cell genes (Fig 6D). This could be indicative of biological differences between single-celled organisms and more complex organisms like maize, or could be a feature of the highly specialized pollen tube. Interestingly, our results are consistent with a recent meta-analysis that identified higher mRNA expression levels as a feature distinguishing gene models with known phenotypes from the overall population of gene models defined by sequencing and other molecular approaches [64]. Given that genes – i.e., those gene models with a clear functional role in determining the characteristics of an organism – represent only a subset of total predicted gene models, genome-scale approaches that can associate quantitative phenotypes with specific genes appear important for achieving a more global understanding of genotype-to-phenotype relationships.

## Methods

### Plant materials

Maize inbred line B73 was used for all RNA isolations. Plants were grown in a controlled greenhouse environment (16 hrs light, 8 hrs dark, 80 F day/70 F night) and in the field at the Botany & Plant Pathology Field Lab (Oregon State University, Corvallis, OR) using standard practices. Lines containing *Ds-GFP* insertion alleles were acquired from the Maize Genetics Cooperation Stock Center.

### RNA isolation, library preparation and sequencing

Detailed methods are available in S1 Methods. Briefly, tissue was isolated either by dissection (TP), differential density centrifugation (MS, MS-B and SC), or collection at anthesis (MP). Total RNA from TP, MS, and MP was extracted using a modified Trizol Reagent (Life Technologies) protocol; SC total RNA was extracted via a phenol/chloroform protocol. Poly-A RNA (mRNA) was isolated using streptavidin magnetic beads (New England Biolabs, # S1420S) and a biotin-linked poly-T primer. RNA libraries were prepared and sequenced by the Central Services Lab (CSL) at the Center for Genome Research and Biocomputing (CGRB, Oregon State University) using WaferGen robotic strand specific RNA preparation (WaferGen Biosystems) with an Illumina TruSeq RNA LT (single index) prep kit and run on an Illumina HiSeq 3000 with 100 bp paired-end reads.

### Mapping reads to genes, differential expression assessment and GO enrichment analysis

Ribosomal reads (rRNA) were removed from all samples using STAR, version 2.5.1b [65] to map reads to a repository of maize rRNA sequences (parameters: --outSAMunmapped Within --outSAMattributes NH HI AS NM MD --outSAMstrandFieldintronMotif --limitBAMsortRAM 50000000000 --outReadsUnmapped Fastx). The number of mappable reads generated from each sample after rRNA removal ranged from approximately 1 million to approximately 41 million, with an average mappable reads of approximately 18 million per sample. Total reads, mapped reads, rRNA contamination, and other statistics are summarized in S1 Table.

rRNA-filtered sequences were mapped to the maize reference genome, version B73 RefGen_v4.33 [39] using STAR, keeping only unique alignments (parameters: -- outSAMunmapped Within --outSAMattributes NH HI AS NM MD --outSAMstrandField intronMotif --outFilterMultimapNmax 1 --limitBAMsortRAM 50000000000). Transcript levels of annotated gene isoforms were measured using Cufflinks, version 2.2.1 [66]. FPKM (fragments per kilobase of transcript per million mapped reads) values are shown in S4 Table. Differential expression was calculated between each tissue with Cuffdiff, version 1.0.2, using default parameters. FPKM counts were normalized using the geometric library normalization method. A pooled dispersion method was used by Cuffdiff to model variance. Differential expression results are summarized in S11 Table.

Gene ontology (GO) terms were found for enriched genes in each tissue using the AgriGO 2: GO Analysis Toolkit [67]. Enriched genes were defined as the top 300 significantly differentially expressed genes (q-value) from Cuffdiff output, with ties broken by log_2_ fold change. Enriched sets were split into up- and down-expressed genes. GO term enrichment was calculated using the singular enrichment analysis method with a Fisher test and Yekutieli multi-test adjustment. GO annotations were based off the maize-GAMER annotation set [68].

### Mapping reads to transposable elements

The rRNA-filtered reads were quality trimmed (QC30) and adapter sequences were removed using BBDUK2 [69]. The remaining sequences were mapped to the whole genome using STAR, allowing mapping to at most 100 ‘best’ matching loci. (parameters: --outMultimapperOrder Random --outSAMmultNmax -1 --outFilterMultimapNmax 100). For paired-end reads, the unmapped reads were re-mapped using single-end approach to maximize the number of mappable reads. The mapping percentage is reported in S3 Table. Because 19% of the total reads in the dataset mapped to more than one location, such reads were mapped to only their best match in the genome, and when multiple best matches existed, they were mapped to all of these loci, and then counted fractionally. For example, if one read maps to 4 TE locations equally well, each TE would receive a weighted value of 0.25 mapped reads. Because the TE expression of the aberrant SC1 biological replicate did not cluster with the other three SC replicates (S1 Fig), it was removed from all subsequent analysis of TE expression.

### Principal component analysis (PCA)

Using the maize gene and TE annotation file available from Ensembl Genomes (v38) [39], a combined annotation file was generated for both genes and TEs to run PCA for all samples. FeatureCounts [70] was used to calculate the accumulation of each gene and TE in all samples following fractional assignment of reads (parameters: -O --largestOverlap -M --fraction -p -C). This counts file was used in DESeq-2 [71] to generate the PCA plot.

### Analysis of transposable elements

From the featurecount file (described above), counts of TEs (farther than 2kb from genes) were retained for further analysis. First, normalized read counts for all TEs were obtained (data in S2 Fig) and then, after selecting seedling as the reference tissue, pairwise volcano plots were generated for all samples against the reference seedling tissue. The number of TEs statistically significantly up- and down-regulated in each tissue was calculated and plotted (Fig 3).

All TEs less than 2kb away from a gene were categorized as ‘near genes’ TEs. Any TE with low expression that was excluded by DESeq for pairwise comparison, was counted in ‘not covered’ category. Compared to seedling, if a TE was found to be up-regulated or down-regulated in any tissue, it was categorized as a dynamic TE. Remaining TEs with p-value > 0.05 compared to seedling were categorized as static TEs, as no evidence of TE expression was observed over different developmental time points analyzed. For all categories, the length, family or distance from centromere was calculated based on the published TE annotation file.

### Validation of *Ds-GFP* insertion sites

A FASTA file containing 2 kb of genomic sequence surrounding each *Ds-GFP* insertion site was used as input to a primer3-based tool to generate a pair of specific primers to genotype individual plants from each line (https://vollbrechtlab.gdcb.iastate.edu/tools/primer-server/). The primers used for each *Ds-GFP* line are listed in S6 Table.

To genotype the plants, two 7 mm discs of leaf tissue were collected from each plant using a modified paper punch. The samples were collected in 1.2 ml tubes that fit within a labeled 96 well plate/rack (https://vollbrechtlab.gdcb.iastate.edu/tools/tissue-sample-plate-mapper/) (Phenix Research Products, Candler, NC; M845 and M845BR or equivalent). Genomic DNA was isolated from the leaf punches [72] with the following modifications. An additional centrifugation (3,000 *g* for 10 min.) was added to clear the leaf extracts prior to loading onto a 96-well glass fiber filter plate (Pall, 8032). DNA was eluted from filter plates in 125 μL water, and 2 μL was used as template for PCR. Amplification followed standard PCR conditions using GoTaq Green Master Mix (Promega) with 4% DMSO (v/v) and amplicons were resolved using agarose gel electrophoresis. Lines were genotyped using the pair of *Ds-GFP* line gene-specific primers plus one *Ds*-specific primer (JSR01 GTTCGAAATCGATCGGGATA or JGP3 ACCCGACCGGATCGTATCGG). All lines were also screened by PCR for the presence of *wx1-m7::Ac* using primers for *wx1* (CACAGCACGTTGCGGATTTC) and *Ac* (CCGGATCGTATCGGTTTTCG). Followup PCR to test for co-segregation of GFP fluorescence with the presence of the insertion used the appropriate set of three PCR primers (two gene-specific and one *Ds-*specific) and DNA prepared either from endosperm or seedling leaves [73].

### Insertional mutagenesis transmission quantification and statistics

Heterozygous lines with PCR-validated *Ds-GFP* insertion alleles were planted in the Botany & Plant Pathology Field Lab (Oregon State University, Corvallis, OR). All insertions were in coding sequence (CDS) sites. Heterozygous *Ds-GFP* plants were outcrossed to tester plants (*c1*/*c1 wx1*/*wx1* or *c1*/*c1* genetic background) through both the female and the male, with male pollinations made with a heavy pollen load on extended silks (silks that had been allowed to grow for at least two days following cutback). Following harvest, resulting ears were imaged using a custom rotational scanner in the presence of a blue light source and orange filter for GFP seed illumination [45]. Briefly, videos were captured of rotating ears, which were then processed to generate flat cylindrical projections covering the surface of the ear (for examples, see Figs 5 and 7). Seeds were manually counted using the Cell Counter plugin of the Fiji distribution of ImageJ [74]. Ears showing evidence of more than a single *Ds-GFP* insertion (∼75% GFP transmission) were excluded from further analysis. Seed transmission rates of remaining ears were quantified using a generalized linear model with a logit link function for binomial counts and a quasi-binomial family to correct for overdispersion between parent lines. Non-Mendelian inheritance was assessed with a quasi-likelihood test with p-values corrected for multiple testing using the Benjamini-Hochberg procedure to control the false discovery rate at 0.05. Significant non-Mendelian segregation was defined with an adjusted p-value < 0.05. Proportions of genes with male-specific transmission defects in the Seedling, Sperm Cell, and Vegetative Cell categories were compared using a two-sided Fisher’s exact test, with significance defined as a p-value < 0.05. A two-sided Fisher’s exact test was also used to compare the proportions of male-specific transmission defects in the most highly expressed genes and the less highly expressed in the vegetative cell category. A two-sided test for equality of proportions with continuity correction was used to compare transmission rates in families with partial *Ac* presence. A Git repository containing statistical tests and plotting information for this portion of the study can be found at https://github.com/fowler-lab-osu/maize_gametophyte_transcriptome.

### gex2 sequence analysis and phenotype characterization

Maize *gex2* protein sequence (Zm00001d005781_T002) was retrieved from the Maize Genetics and Genomics Database (MaizeGDB) hosting of the B73 v4 genome [39,75]. Arabidopsis *GEX2* protein sequence (AT5G49150.3) was retrieved from the Arabidopsis Information Portal (ARAPORT) Col-0 Araport11 release [76,77]. Protein sequences were aligned using EMBOSS Needle [78]. Maize and Arabidopsis *gex2* protein domains were predicted by InterPro [79], with transmembrane helix predictions by TMHMM [80]. Prediction of land plant species *gex2* conservation was retrieved from PLAZA, gene family HOM04M006791 [81]. Maize *gex2* gene duplication searches were performed using BLAST [82] and the B73 v4 genome. To confirm the predicted insertion sites for the two *gex2::Ds-GFP* alleles, flanking insertion site fragments were PCR-amplified with a gene-specific primer and a *Ds-GFP-*specific primer (DsGFP_3UTR – TGCAAGCTCGAGTTTCTCCA) and sequenced via Sanger sequencing.

To quantify small seed phenotype, mature, dried down maize ears were imaged prior to seed removal from the ear. For small seeds selection, the ear was first visually scanned row by row from the top to the bottom of the ear. Seeds that were noticeably smaller than their surrounding (regular-sized) seeds are carefully removed from the ear using a pin tool. This sometimes required the removal of regular-sized surrounding seeds, which were saved for later counting. A second visual inspection of the ear often resulted in additional small seeds and is recommended. All remaining seeds were then removed from the ear by hand or using a hand corn sheller tool (Seedburo Equip. Co., Chicago, IL). The ear was screened again for any small (flat/tiny) seeds that could have been missed previously. The cob was inspected prior to discarding, and if any small seed was left behind it was removed and accounted for. Small/smaller seeds and regular-sized seeds were counted and counts were recorded (S10 Table). To measure seedless area, ears were scanned as previously described to create flat surface projections. “Seedless area” was defined as ear surface area that lacked mature or partially developed seeds. Seedless area was quantified as a percentage of total area, as measured with the “Freehand selection” tool of the Fiji distribution of ImageJ [74]. A Git repository containing statistical tests and plotting information for this portion of the study can be found at https://github.com/fowler-lab-osu/maize_gametophyte_transcriptome.

For analysis of embryo sacs by confocal microscopy, tissues were stained with acriflavine, followed by propidium iodide staining [83,84]. After staining, samples were dehydrated in an ethanol series and cleared in methyl salicylate. Samples were visualized on a Leica SP8 point-scanning confocal microscope using excitations of 436 nm and 536 nm and emissions of 540 ± 20 nm and 640 ± 20 nm.

## Acknowledgements

We thank O. Childress, H. Fowler, B. Galardi, B. Hamilton, R. Hartman, and C. Lambert for their tireless seed counting, genotyping, field work, and other technical assistance; and Dr. Lian Zhou for her contributions to maize field genetics. We also thank K. Wimalanathan and T. Shibamoto for computational support at ISU, and M. Dasenko and the Center for Genome Research and Biocomputing for library preparation, sequencing and computational support at OSU. We thank D. Auger for reading the manuscript.

## Supporting information

**S1 Fig. Principal component analysis of gene and transposable element (TE) expression levels**.

Two major components, on x- and y-axis, explain 89% of the variance in gene and TE expression levels. Asterisk (*) mark indicates the sample generated as part of this study, whereas other datasets are publicly available. For the sperm cells isolated in this study (SC), the TE expression of one biological replicate did not cluster with the other three (SC1), and therefore was removed from subsequent analyses of expression from TEs. MP-2014, SE, and OV are from [26]; MP-WEB is from [41]; LF is from [40]; MP-LM is from NCBI BioProject 306885 (2015); SC-TD is from [30].

**S2 Fig. Seedling tissue is the appropriate reference for comparison of TE activity**.

**(A)** Steady-state mRNA accumulation of all TEs in different tissues. Datasets generated in this study are marked with an asterisk. **(B)** The number of TEs with zero or near-zero expression levels in different tissues. Seedlings (SE) have the most TEs with low expression levels.

**S3 Fig. Abundance of TEs near genes in each tissue**.

**(A)** For each tissue type, the top 20,000 expressed genes are distributed along the X-axis in bins of 200, with the highest expressed bin on the far left. The number of TEs near (<2kb) these genes is then counted on the Y-axis. **(B)** Genes filtered for either higher expression in pollen (MP) over sperm cells (SC) (left) or SC>MP (right) were used to determine if the association in Figure 4 is due to sample contamination between SC and MP. Once genes were filtered, the top expressed genes in that tissue were distributed along the X-axis in bins of 200 based on their expression values, with the highest expressed bin on the far left. The number of up- and down-regulated TEs near (<2kb) these genes is then counted on the Y-axis.

**S4 Fig. *gex2* mutant pollen is associated with increased small and aborted seeds in outcross progeny**.

**(A)** Seeds were removed from ears, arranged according to size, and counted. Images of representative seed populations are shown, with the top two rows in each image showing representative fully developed seeds. Rows below the top two contain all of the smaller or aborted seed from that particular ear. **(B)** PCR genotyping of small endosperm seeds from two independent crosses for the two *gex2* alleles show the majority of small seeds harbor the *gex2::Ds-GFP* allele, despite overall reduced transmission of the insertion alleles through the male. Small seeds from control *tsdgR46C04* crosses segregate in a Mendelian fashion.

**S5 Fig. Characterization of gex2 seedless ear area**.

Seedless area was quantified from scanned ear images for *gex2 Ds-GFP* alleles and *Ds-GFP* controls. Pollen from heterozygous *gex2* plants did not show significantly increased seedless area (*gex2-tdsgR82A03* pairwise t-test p-values relative to GFP line 1, GFP line 2, and VC mutant 0.95, 0.96, and 0.74, respectively; *gex2-tdsgR84A12* pairwise t-test p-values 0.19, 0.13, and 0.06, respectively), whereas pollen from homozygous *gex2-tdsgR84A12* plants had significantly increased seedless area (pairwise t-test, p-value < 0.0001).

**S1 Methods. Tissue sample preparation, RNA extraction, and analysis of potential confounding variables in insertional mutagenesis lines.**

**S1 Table. Gene sequencing statistics and availability**.

Summary statistics for sequencing data generated in the study.

**S2 Table. GO term enrichment results**.

Differentially expressed genes in developmental categories examined in the study, as well as significantly enriched GO terms associated with these genes.

**S3 Table. Transposable element sequencing statistics and availability**.

Summary statistics and availability for expression datasets used in the analysis of transposable element expression.

**S4 Table. Genic isoform abundance (FPKM) across developmental stages**.

Cufflinks output describing isoform expression by developmental stages, separated by biological replicate.

**S5 Table. Top 20% transcripts by FPKM in Mature Pollen, Sperm Cell and Seedling datasets**.

List of top 20% highly expressed genes assigned to the Vegetative Cell, Sperm Cell or Seedling Only classes.

**S6 Table. Insertional mutagenesis alleles and primers**.

List of alleles tested for the presence of *Ds-GFP* insertions by PCR, including primers sequences.

**S7 Table. Insertional mutagenesis results by line**.

Insertional mutagenesis results, separated by line, including marker transmission rates and expression category information.

**S8 Table. Insertional mutagenesis results by allele**.

Insertional mutagenesis results, separated by allele, including marker transmission rates and expression category information.

**S9 Table. Concordance of seed phenotype with DsGFP genotype**.

PCR results from testing *Ds-GFP* presence GFP and non-GFP seeds for selected alleles.

**S10 Table. gex2 small seed phenotyping**.

*gex2* small seed counting and seedless area results.

**S11 Table. Differential expression results**.

Cuffdiff output comparing expression between tissues examined in this study.

